# Assembly Defects Abrogate Proofreading by Initiation Factors and License the Entry of Premature Ribosomes into the Translation Cycle

**DOI:** 10.1101/498592

**Authors:** Himanshu Sharma, B. Anand

**Affiliations:** Department of Biosciences and Bioengineering, Indian Institute of Technology Guwahati, Guwahati 781039, Assam, India

**Keywords:** Ribosome, Ribosome assembly, Assembly factors, Assembly defects, Translation initiation, Translation fidelity, Decoding, Initiation factors

## Abstract

Numerous quality control steps are deployed by the translational machinery to ensure faithful decoding of genetic message to synthesize proteins. However, what transpires to quality control mechanism during protein synthesis when the ribosomes are produced with assembly defects remains enigmatic. In *E. coli*, we show that ribosomes with assembly defects evade the proofreading steps during translation initiation and participate in the translation cycle. Such ribosomes show severely compromised decoding capabilities that give rise to errors in initiation and elongation. Tracing the genesis, we discovered that the assembly defects compromise the binding of initiation factors, thus licensing the rapid transitioning of 30S (pre) initiation complex to 70S initiation complex by tempering the proofreading mechanism. Overall, our work highlights that a mass balance deficit between premature ribosomes and initiation factors steers the entry of premature ribosomes into the translation cycle.

## Introduction

The bacterial ribosome is a large ribonucleoprotein complex consisting of two asymmetrical subunits 30S and 50S, respectively. The subunits are composed of three ribosomal RNAs (5S, 16S & 23S) and more than 50 ribosomal proteins (r-Proteins). Ribosomes are synthesized by a highly regulated process, which is collectively referred as ribosome biogenesis that involves the synthesis of their components and their concomitant association. The orchestrated association of r-Proteins with rRNA – termed ribosome assembly – is catalysed by some non-ribosomal proteins called as ribosome assembly factors (RAF) (Davis and Williamson, 2017; Shajani et al., 2011). RAFs are involved in end processing of the premature rRNA (Baumgardt et al., 2018; Deutscher, 2015; Smith et al., 2018; Sulthana and Deutscher, 2013) and rRNA modifications like pseudouridylation (Ψ) and methylation (Boehringer et al., 2012; Jack et al., 2011; Polikanov et al., 2015; Spenkuch et al., 2014). Additionally, RAFs assist in rRNA folding and positioning of r-Proteins thus overcoming energy barriers and kinetic traps during assembly (Charollais et al., 2004; Charollais et al., 2003; Gulati et al., 2014). Finally, RAFs also mark important maturation events during assembly (Goto et al., 2011) thus driving the creation of functional ribosomal subunits.

The subunits that endure full maturation become competent to enter the translation cycle. The labour of translation is divided between the two subunits: the 30S subunit decodes the genetic message and the 50S subunit catalyses the peptide bond formation. The 30S subunit interacts with mRNA, tRNA and initiation factors (IFs) to form the pre-Initiation complex (30S-PIC). During this process, the IFs perform critical quality control tests on codon-anticodon (CO-AC) triplets on mRNA and tRNA, respectively, and this leads to the formation of 30S-IC (Gualerzi and Pon, 2015; Hussain et al., 2016; Simonetti et al., 2008). 30S-IC associates with 50S subunit to form the 70S initiation complex (70S-IC). The 70S-IC eventually enters the elongation phase of the translation and drives protein synthesis. Disruptions in translation are known to hinder the biogenesis of ribosomes by skewing ration of r-Proteins to r-RNA (Nikolay et al., 2014; Siibak et al., 2009). Additionally, the suboptimal maturation of ribosomes is implicated in several disorders collectively referred to as ribosomopathies (Danilova and Gazda, 2015; Freed et al., 2010; Mills and Green, 2017). However, the functional consequences of assembly defects on translation are yet to be fully understood.

Assembly defects arising due to deletion of RAFs are manifested as compromised growth phenotype, sensitivity to cold temperatures as well as accumulation of ribosome assembly intermediates (Alix and Guerin, 1993; Boehringer et al., 2012; Britton, 2009; Bylund et al., 1998; Charollais et al., 2004; Charollais et al., 2003; Davis and Williamson, 2017; Guthrie et al., 1969; Shajani et al., 2011). These intermediates harbour premature rRNA as well as a suboptimal r-Protein complement. Efforts to understand the nature of structural defects in assembly intermediates in bacteria are undermined by pleiotropy associated with deletion of the RAFs and the presence of multiple pathways for the maturation of ribosomes (Earnest et al., 2015; Lai et al., 2013; Lerner and Inouye, 1991; Talkington et al., 2005). Additionally, little is known about the metabolic cost of harbouring premature subunits and their ultimate fate in bacterial cells. Although studies have shed light on how cells regulate the number of active ribosomes and maintain fitness in nutrient-limiting conditions (Dai et al., 2018; Dai et al., 2017; Koch and Deppe, 1971; Neidhardt and Magasanik, 1960; Towbin et al., 2017), such knowledge is virtually unknown for cells with compromised ribosome structure and composition.

It is believed that subunit association may be impaired in premature ribosomes, which lack fully formed bridging sites, and this, in turn, may arrest their entry into translation (Boehringer et al., 2012; Datta et al., 2007; Razi et al., 2017). In contrast, it has also been observed that mutations in rRNA, lack of r-Proteins or the absence of assembly factors like KsgA and Hfq induce defects during translation (Andrade et al., 2018; Connolly and Culver, 2013; O’Connor et al., 2004; Qin et al., 2012). Studies have also hypothesized that perturbed ribosome maturation may create a bottleneck that decreases new translation initiation events (Gibbs et al., 2017). This would result in the suppression of translation or alternate scenarios wherein premature subunits may compete with mature subunits for the mRNA thus averting translation once again. The spectrum of studies suggests that participation of premature ribosomes in translation may have energetic implications that affect the cellular fitness. However, mechanistic details of such relationships are yet to be uncovered.

Here, we set out to address some of the primary outstanding questions: Do premature ribosomes enter translation? If so, how such ribosomes bypass the quality control mechanisms in bacteria? To address these questions, we employed 30S from *Escherichia coli* as a model system and assessed how the defects in the 30S assembly engender errors in translation initiation and elongation. Using a series of biochemical and next-generation sequencing experiments, we establish that premature ribosomes indeed enter the translation cycle but with compromised fidelity in decoding. Our investigation led us to establish that the assembly defects weaken the recognition by the IFs, especially IF3, and thereby permits this bypass of quality control mechanisms.

## Results

### Defects in late 30S assembly disrupt fidelity of translation

We wondered whether the premature 30S subunits bypass the quality control checkpoints to enter the translation cycle. To address this, we created several variants of *E. coli* that harboured null mutants for genes encoding late-stage 30S specific RAFs such as RsgA, RbfA, KsgA and LepA (Fig. 1A). Premature ribosomes from these strains were tested for their ability to recognize AUG (cognate) and non-AUG (near cognate and non-cognate) initiation codons for translating the gene encoding GFP *in vivo* (Fig. 1B). In order to quantify the extent of error in decoding the start codon due to an assembly defect in the 30S subunit, we formulated two indices: (i) Initiation Error Index (IEI) and (ii) Initiation Rescue Index (IRI). IEI was defined as the ratio of GFP fluorescence from null mutants of the respective RAF to Wild type (Wt). We have chosen four different initiation signals of variable strength: AUG>CUG>AUA>AGG to assess the potency of premature 30S subunits in discriminating cognate from non-cognate codons (Hecht et al., 2017; Sussman et al., 1996).

**Figure 1.**
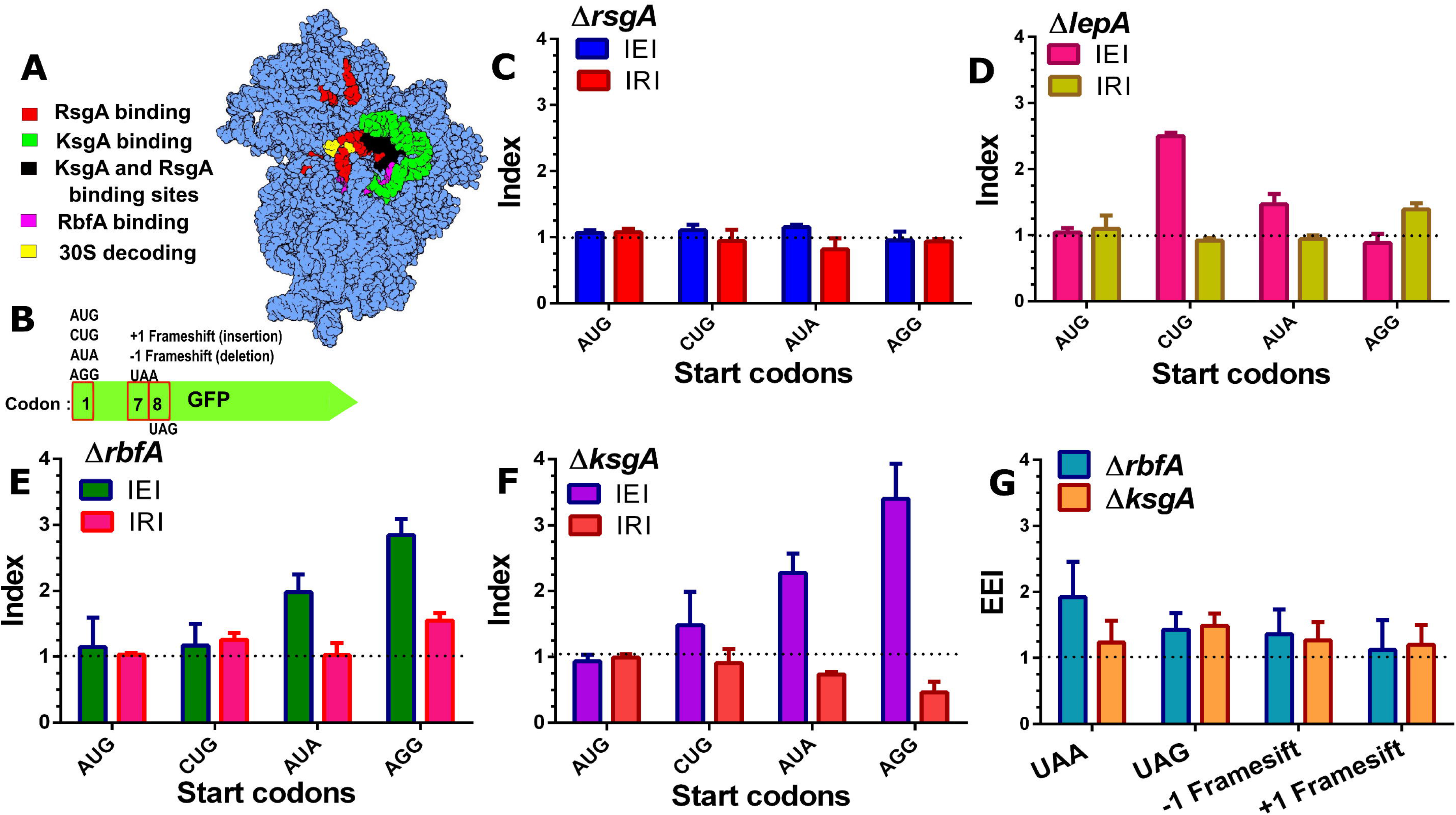
Assembly defects in 30S subunit compromise decoding fidelity. **(A)** Structure of the *E. coli* 30S subunit represented in blue spheres [PDB ID: 4V52] showing the binding sites of late stage acting RAFs as observed by Cryo-EM studies (Boehringer et al., 2012; Datta et al., 2007; Razi et al., 2017). Colour coding for each RAF binding site is indicated along with the 30S decoding centre. **(B)** A representation of the GFP construct used for measuring translation fidelity. Codon variations at codon 1, 7 and 8 are indicated. **(C)– (F)** Errors and rescue from misrecognition of start codon, measured as IEI and IRI are shown for *ΔrsgA* (C), *ΔlepA* (D), *ΔrbfA* (E) and *ΔksgA* (F), respectively. Dashed lines marking IEI =1, signifies similar expression in null mutants and Wt, indicating optimal recognition of the start codon. Dashed lines marking IRI =1, signifies rescue from erroneous translation initiation at near or non-canonical start codons. **(G)** Evaluation of frame maintenance and fidelity of stop codon recognition measured as EEI, is shown for *ΔksgA* and *ΔrbfA*. The dashed line indicating EEI =1 indicates similar level of GFP expression between null mutants and Wt cells.

Deletion of RsgA did not display any increase in the IEI, indicating no difference in the codon recognition between *ΔrsgA* and Wt (Fig. 1C). Deletion of the recently implicated RAF, EF-4/LepA led to elevated misrecognition of CUG and AUA as start codons (Fig. 1D). However, expression from AUG and AGG start codons remained unchanged in *ΔlepA* strain (Fig. 1D). In order to further investigate if the decoding errors were specifically due to deletion of the respective assembly factor, we complemented the loss of RsgA and LepA from a plasmid-borne copy. Initiation Rescue Index (IRI) was calculated as the ratio of GFP fluorescence in complemented null mutant to that of Wt. If the translation errors were indeed caused due to the absence of the respective RAF, a shift from IEI >1 to IRI ∼1 would hint at a rescue from misinitiation at non-canonical start codon. Upon complementation, IRI∼1 indicated that translation defects were ameliorated in *ΔlepA* cells (Fig.1D), on the other hand, the IRI remained unaffected for *ΔrsgA* cells (Fig. 1C). In contrast, loss of RbfA and KsgA displayed an elevated IEI for near-cognate as well as non-cognate start codons indicating significant misinitiation (Fig. 1E & 1F). In order to probe this phenomenon further, we also complemented the loss of RbfA and KsgA with a plasmid-borne copy to measure the IRI values. The rescue was indicated by IRI ∼ 1 in RbfA and KsgA complemented strains (Fig. 1E & 1F). These findings highlight the fact that the premature ribosomes accumulating due to loss of RsgA, LepA, KsgA and RbfA have the potency to enter translation even with a significantly compromised native structure.

Given the aberrant effect on the initiation stage exerted by the premature ribosomes, we tried to investigate further if the assembly defects in *ΔrbfA* and *ΔksgA* strains also manifest during translation elongation stage. Towards this, we employed variants of *gfp* that harboured frameshift mutations or premature stop codons and measured the Elongation Error Index (EEI) (Fig. 1G). EEI >1 would indicate decoding errors leading to compromised frame maintenance or bypass of stop codons with respect to Wt. Both *ΔrbfA* and *ΔksgA* showed slightly elevated EEI values, indicating that premature ribosomes in both strains have compromised fidelity during elongation and termination. However, the observations also suggest that these bypasses are less frequent in comparison to the erroneous decoding of the initiation signals.

### Defects in assembly upset the kinetics of translation initiation and elongation

In order to further dissect the link between assembly and translation, we studied translation by tracing the pre-steady-state kinetics of β-galactosidase (bgal) synthesis *in vivo*. For this, we employed an IPTG inducible construct, pAUG-bgal (AUG as the start codon). The trace signifying accumulation of bgal was used to calculate the peptide chain elongation rate (ER) as described previously (Dalbow and Young, 1975; Schleif et al., 1973). We measured ER for Wt, *ΔrbfA* and *ΔksgA* at 37°C (Fig. 2A; Table 1). Wt was found to have ER that was in close agreement with the previous report (Dai et al., 2017). However, the elongation was slightly faster in case of *ΔksgA*, whereas it was slowed down by approximately three-fold for *ΔrbfA* relative to Wt (Fig. 2A & 2C; Table 1), suggesting that the perturbed assembly indeed upsets translation. This was even more striking at 25°C and was in line with the assembly defects getting exacerbated at lower temperature (Fig. 2B & 2D; Table 1).

**Figure 2.**
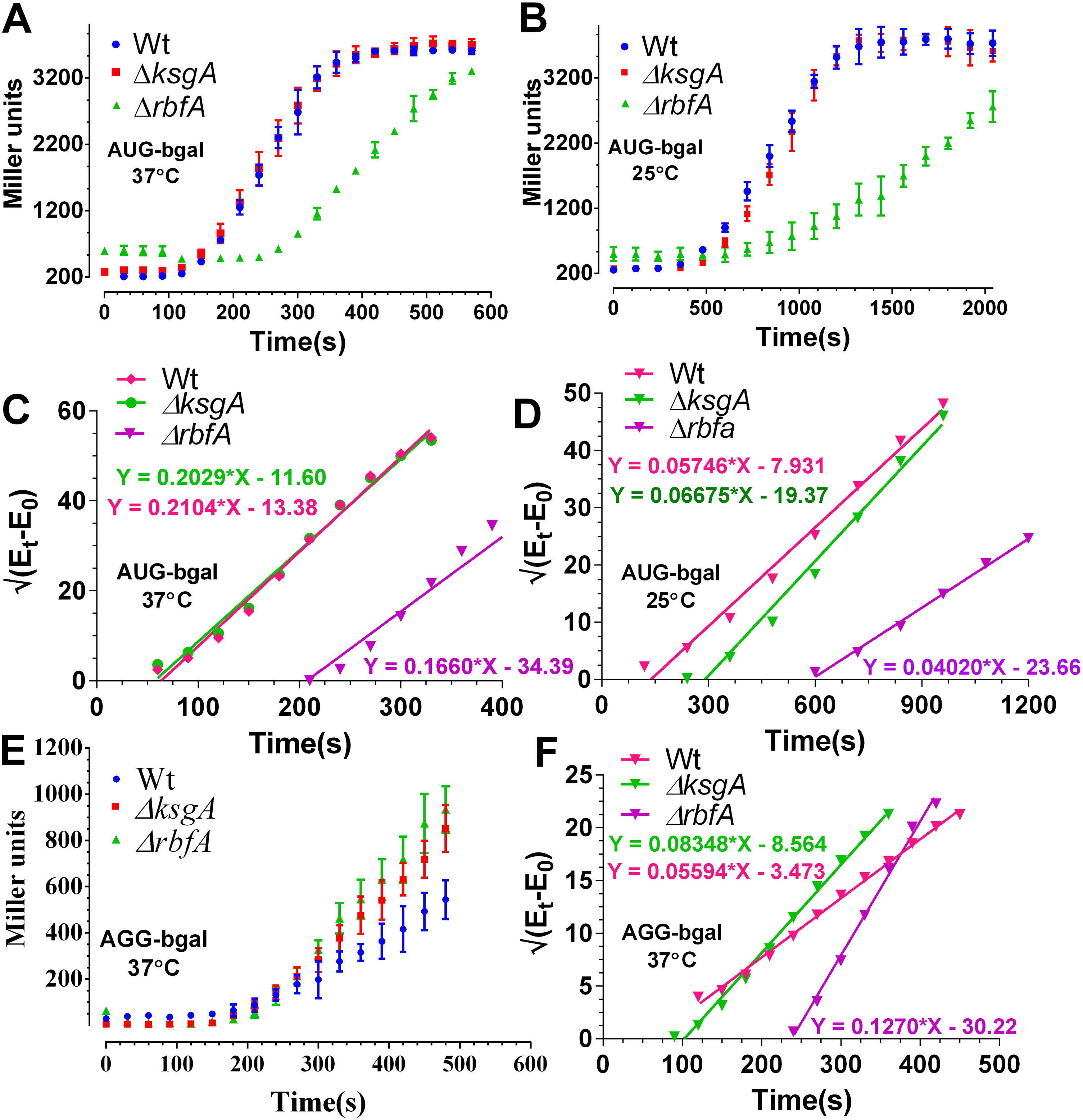
Translation kinetics of premature ribosomes. **(A)–(B)** Translation kinetics of bgal production with AUG start codon in Wt, *ΔrbfA* and *ΔksgA*. Protein production kinetics was performed at 37 °C (A) and 25 °C (B), respectively. The plots were drawn as averaged measurement from three independent time course experiments. The error bars represent standard deviation of the three independent trials. **(C)-(D)** Schleif plot for bgal translation kinetics derived by plotting the square root of residual bgal enzyme activity against time from data in (A) & (B) (*vide*. Methods). The plotted data was fitted to a linear regression model. Equations representing the trend line for each plot have been shown. The slope of the line indicates the rate of translation initiation and the X-intercept marks the first appearance of the enzyme activity that can be further used to calculate the rate of peptide chain elongation. **(E)** Translation kinetics of bgal production with AGG start codon in Wt, *ΔrbfA* and *ΔksgA*. The kinetics was performed at 37 °C. **(F)** Schleif plot for bgal translation kinetics with AGG start codon derived from the data in (E).

**Table 1:** Peptide chain elongation rate (ER) measured from Wt, *ΔksgA*, and *ΔrbfA c*ells using bgal translation kinetics. All measurements are in amino acids/second (aa/sec).

| Start codon | Growth temperature | Wt | $\Delta ksgA$ | $\Delta rbfA$ |
| --- | --- | --- | --- | --- |
| AUG | 37 °C | 16.1 ± 2.5 | 17.9 ± 2.9 | 4.9 ± 0.5 |
| AUG | 25 °C | 7.4 ± 1.9 | 3.5 ± 0.5 | 1.7 ± 0.1 |
| AGG | 37 °C | 16.1 ± 3 | 9.94 ± 0.7 | 4.3 ± 0.2 |
| AGG | 25 °C | -ND- | -ND- | -ND- |

Using the same kinetics data, we also tried to understand how the assembly defects affect the translation initiation process. We measured the translation initiation rate (IR), as indicated by the slope of the bgal accumulation trace (Fig. 2C & 2D) (Dalbow and Young, 1975; Schleif et al., 1973). The rate of translation initiation from the AUG start codon followed a pattern similar to the ER. Rate of initiation was similar for Wt and *ΔksgA* (Fig. 2C; Table 2) and it was lower for *ΔrbfA* with respect to WT at 37°C (Fig. 2C; Table 2). In line with ER, IR for Wt, *ΔrbfA* and *ΔksgA* was significantly reduced at 25°C (Fig. 2D; Table 2). Notably, IR for *ΔrbfA* was consistently lower relative to WT, reinforcing that premature ribosomes entering translation in *ΔrbfA* have impaired translation capacity.

**Table 2:** Translation initiation rates (IR) measured from Wt, *ΔksgA*, and *ΔrbfA* cells using bgal translation kinetics. The IR values are derived from the slope of the curve marking accumulation of bgal over time (units: Miller units^1/2^/sec).

| Start codon | Growth temperature | Wt | $\Delta ksgA$ | $\Delta rbfA$ |
| --- | --- | --- | --- | --- |
| AUG | 37 °C | 0.2104 ± 0.008 | 0.2029 ± 0.007 | 0.1660 ± 0.008 |
| AUG | 25 °C | 0.05746 ± 0.002 | 0.06675 ± 0.003 | 0.04020 ± 0.001 |
| AGG | 37 °C | 0.05594 ± 0.001 | 0.08348 ± 0.002 | 0.1270 ± 0.004 |
| AGG | 25 °C | -ND- | -ND- | -ND- |

It was also important to address a possibility whether the aberrant ER and IR represented errors in decoding or just a general slowdown of the translational machinery that might arise due to a deficit in number of mature ribosomes entering translation. In order to address this, we hypothesized that, if the aberrant ER and IR were caused by a general slowdown of the translational machinery, the respective rates should remain unaffected if measured from non-canonical start codons. However, if the decrease was indeed caused by erroneous decoding due to assembly defects, the non-canonical start codons must be misread as canonical ones preferentially in null mutants in comparison to Wt. To test this notion, we performed bgal translation kinetics using constructs with an AGG start codon (pAGG-bgal) at 37°C. Remarkably, the IR from AGG start codon displayed an inverse pattern in comparison to AUG start codon for the three strains. Null mutants of RbfA displayed elevated translation initiation over time in comparison to Wt (Fig. 2E & 2F; Table 2). Similar reversal, but with a lesser intensity, was also observed for *ΔksgA* (Fig. 2E & 2F; Table 2). Unexpectedly, the ER from AGG start codon for Wt and *ΔrbfA* remained unaffected, whereas the elongation was drastically slowed down for *ΔksgA* (Fig. 2F; Table 1). These observations strongly support the conjecture that the errors in assembly lead to erroneous translation initiation and elongation events. Collectively, these findings hint at a scenario where evasion of quality control checkpoints permits entry of premature ribosomes into translation. Such ribosomes not only attenuate the intensity of the translation process but also elude the quality control checkpoints during translation initiation.

### Deep sequencing based profiling of premature ribosomes reveals genome-wide alteration of ribosome occupancy during translation initiation

In order to probe how the assembly defects impact the formation of initiation complexes at genomic level, we turned to TCP-seq (Archer et al., 2016; Shirokikh et al., 2017), a modified Ribo-seq strategy that has been devised to probe translation initiation, elongation and termination complexes in *Saccharomyces cerevisiae* (Fig. 3A). For bacterial Ribo-seq, widely used bacterial translation elongation inhibitors like Chloramphenicol do not trap ribosomes in the initiation stage and also introduces a codon specific bias when used to arrest ribosomes during elongation (Marks et al., 2016). In contrast, TCP-seq uses formaldehyde cross-linking to trap ribosomes during various stages of translation *in vivo* and we have adopted this technique for probing the translational complexes in *E. coli*. In order to arrest the ribosome from Wt, *ΔksgA* and *ΔrbfA* (Fig. 3B) during various stages of translation in an unbiased manner, we snap chilled the exponentially growing cells and cross-linked mRNA bound translating complexes using formaldehyde (Archer et al., 2016; Valášek et al., 2007). After cross-linking, free 30S particles were depleted using sedimentation analysis and mRNA cross-linked ribosomes were released by treating with Micrococcal nuclease (MNase). The released particles were further partitioned using density gradient sedimentation and then used to extract the mRNA footprints (FP) bound to 30S and 70S particles. For library preparation, a wide range of FPs of roughly 10–80 nucleotides (nt) were size selected to accommodate FPs arising from translating complexes as well as the tRNA molecules.

**Figure 3:**
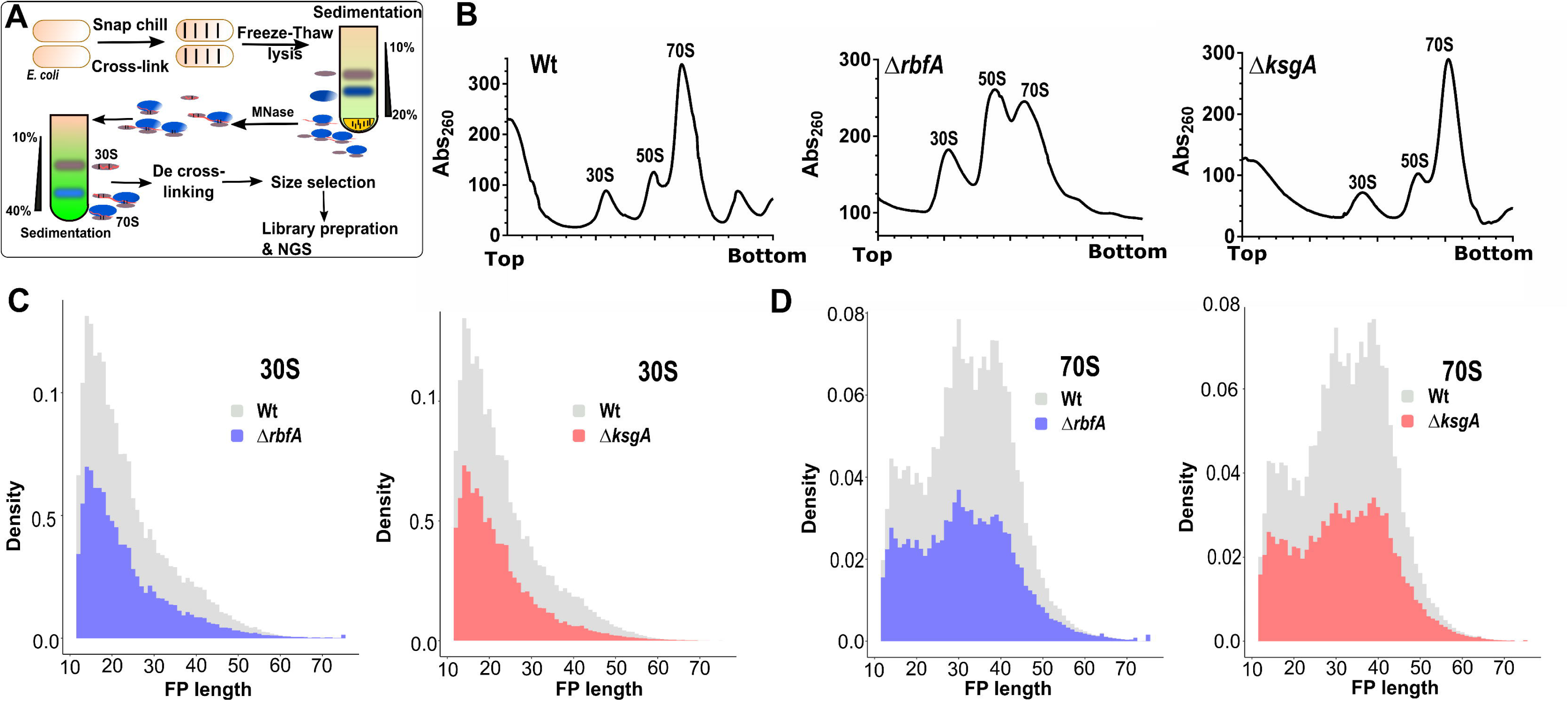
Translation complex profiling for studying initiation and elongation. **(A)** An outline of the strategy used to capture the ribosomes at various stages of translation using TCP-seq. **(B)** Representative polysome profiles for Wt, *ΔrbfA*, and *ΔksgA* derived by performing sucrose density gradient sedimentation. Samples for TCP-seq were derived from 30S and 70S fractions, respectively. **(C)** The distribution of the length of 30S protected fragments isolated from Wt, *ΔrbfA* and *ΔksgA* is shown. **(D)** The distribution of the length of 70S protected fragments isolated from Wt, *ΔrbfA* and *ΔksgA* is shown.

The length distribution of FPs that were mapped to the coding region was congruent with the size selection criteria used during library preparation, harbouring reads ranging from as low as 12 nt through 75 nt in length (Fig 3C & 3D). The 30S subunit derived FPs (30S_FP) contained a distinct population of ∼ 14 nt in length (Fig. 3C). Whereas, we spotted three distinct population of FPs from 70S particles (70S_FP) centred on ∼ 15, 30 and 40 nt, respectively (Fig. 3D). Deletion of both KsgA and RbfA led to a significant drop in the number of FPs, suggesting altered translation with respect to Wt (Fig. 3C & 3D). The size of the FPs mirrors the conformational rearrangements the ribosomes undergo as well as their interactions with translation factors during different stages of translation (Archer et al., 2016; Lareau et al., 2014). To track these, we mapped the 5’ and 3’ end of the FPs of given sizes to the known START of the 4257 coding sequences (CDS) in *E. coli* (Fig. 4). In line with the characteristic size distribution of FPs for 30S and 70S, the majority of the mapped FPs for 30S fell in the smaller size regime (12 – 20 nt) whereas they corresponded to larger size regime (30 nt and above) for 70S (Fig. 4). Each FP of a given length showed a range of characteristic offset distances from the start codons. 30S_FPs were densely populated near the start codon as against 70S_FPs that recapitulated the fact that 30S is subjected to extensive regulatory events during initiation. At the 5’ UTR, the trailing edge of 30S_FP (5’ end of FPs in Fig. 4A) showed graded increase in the offset distance with increase in the length of FPs than the leading edge (3’ end of FPs in Fig. 4C). In fact, the leading edge was arrested around the start codon, suggesting that there is a queuing of 30S ribosomes at the 5’ UTR due to the fact that the residence time of 30S at the start codon is protracted owing to the extensive regulatory controls. However, these signatures were significantly altered in case of *ΔksgA* and *ΔrbfA* especially for 30S than for 70S. This suggests two possibilities: Either the premature ribosomes are actively rejected by the translational machinery to enter the translation cycle or these ribosomes are rapidly surpassing the elaborate checkpoints during the initiation stage (Fig. 4). Both of these cases will reduce the residence time of ribosomes around the start codon and thereby the ribosome occupancy. However, in line with the translational kinetics studied using bgal assay that showed minor reduction in translation initiation for the premature ribosomes (Fig. 2, Table 1 & 2), we reasoned that the altered ribosome occupancy for *ΔksgA* and *ΔrbfA* could be attributed to rapid surpassing of the kinetic checkpoints during the initiation.

**Figure 4:**
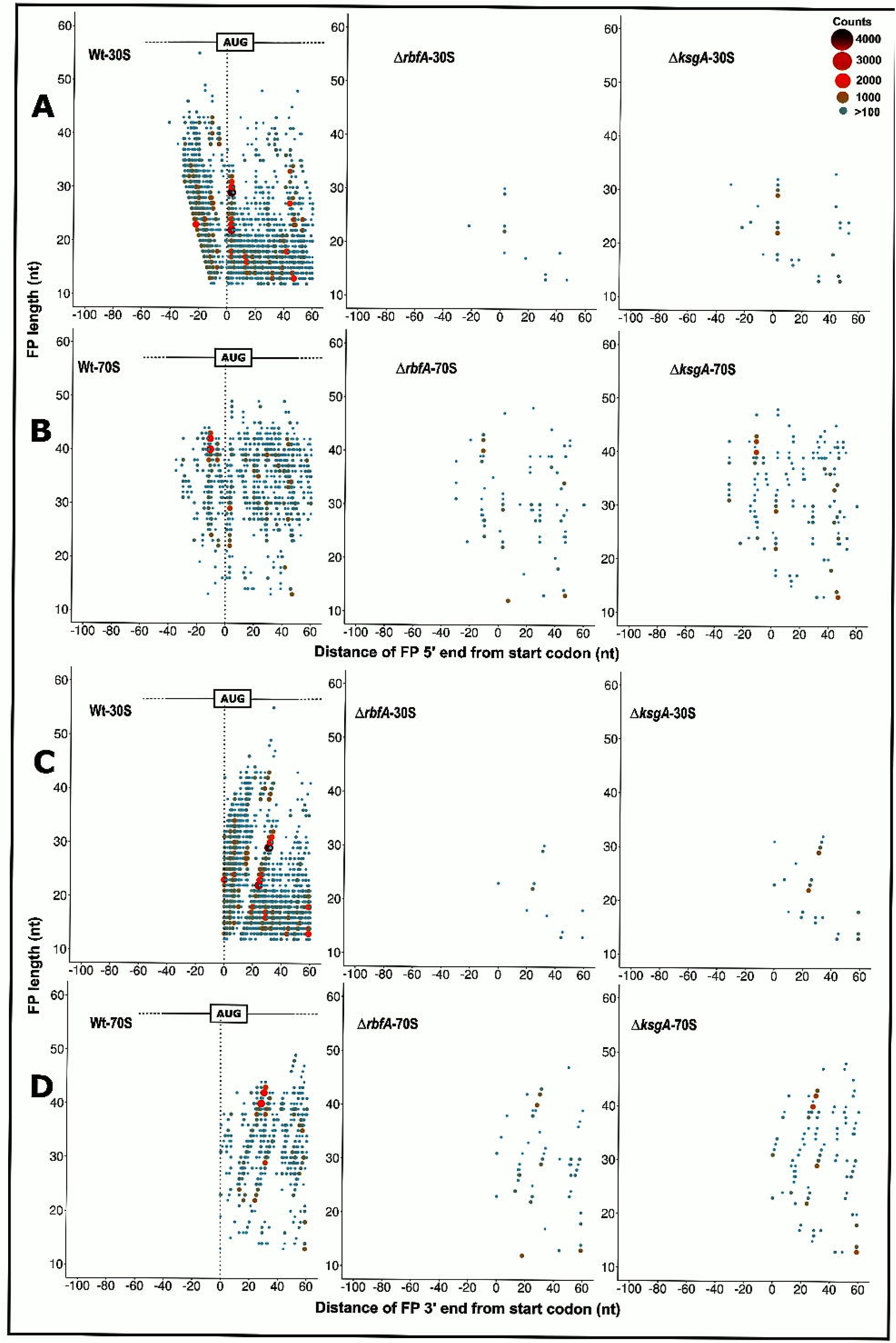
Metagene analysis for translation initiation complex. 5’ ends of the FPs derived from 30S **(A)** and 70S **(B)** bound mRNA as well as the 3’ ends of the FPs derived from 30S **(C)** and 70S **(D)** from Wt, *ΔrbfA* and, *ΔksgA*, respectively, are mapped against the known start codon of the coding sequences in *E. coli*. The first base of the start codon is aligned to the ribosome P site. The size and colour of the points correspond to the frequency of occurrence of FPs. Only those FPs with at least 100 reads are represented here.

### Premature 30S forms short-lived preinitiation complexes

In order to further understand the nature of initiation and elongation complexes formed by the premature subunits, we studied the distribution of N-Formylmethionine initiator–tRNA (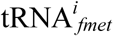) and the elongator tRNA (tRNA*^e^*) that cross-linked and co-purified with the 30S and 70S fractions (Fig. 3A). We reasoned that the tRNA distribution (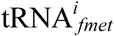 and tRNA*^e^*) would act as an indicator to pinpoint the ribosomes that are at initiation and elongation stages of the translation. The initiation stage being the rate-limiting step of translation is expected to be overpopulated by 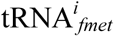 during the transition of 30S-PIC to 30S-IC (Archer et al., 2016; Milon et al., 2012) as compared to other stages associated with the elongating 70S particles. Similarly, an inverse distribution of tRNA*^e^* is expected for elongating 70S than the 30S-PIC/IC. In line with our hypothesis, Wt 30S ribosomes were found to harbour more number of 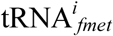 than the 70S (Fig. 5A). Surprisingly, the distribution of 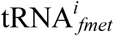 was reversed in both *ΔksgA* and *ΔrbfA* such that its level in 70S was significantly higher in comparison to that in 30S. We posit that the reduced distribution of 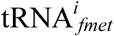 in the 30S fraction of *ΔksgA* and *ΔrbfA* may arise from diminished binding affinity of pre-30S towards the 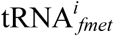 or due to a short lived pre-30S-PIC/IC formed in *ΔksgA* and *ΔrbfA*, which rapidly associate with the 50S subunit to form 70S-IC. Since the transition from 30S-PIC/IC to 70S IC occurs through a well-orchestrated sequence of events, the association between 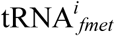 and 70S should have occurred during the formation of 70S-IC. Therefore, this alludes to the possibility that there is a rapid transition of pre-30S-PIC/IC to 70S-IC in *ΔksgA* and *ΔrbfA* and the prospect of low affinity between pre-30S and 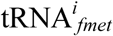 can thus be ruled out. This is further bolstered by the altered ribosome occupancy observed for *ΔksgA* and *ΔrbfA* during metagene analysis (Fig. 4). Further, tRNA*^e^* was distributed approximately in equal measure between Wt 30S and 70S particles, suggesting the dynamic interaction between tRNA*^e^* and ribosome subunits. However, its distribution was skewed for *ΔksgA* and *ΔrbfA* and showed more preference for 70S than the 30S (Fig. 5A).

**Figure 5:**
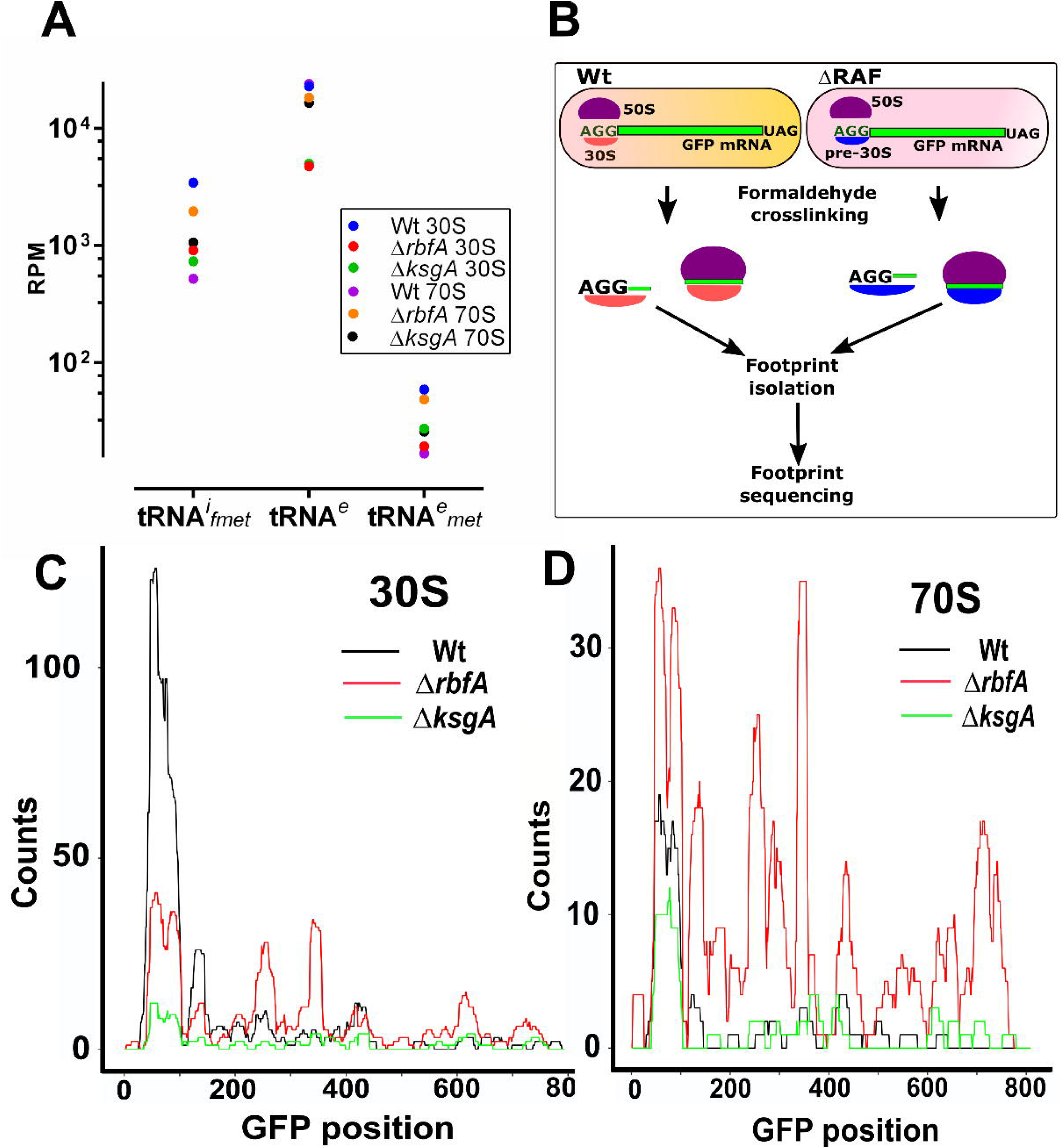
Premature ribosomes form distinct initiation complexes. **(A)** Distribution of initiator (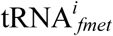) and elongator (tRNA*^e^*) tRNAs in 30S and 70S derived fractions as observed in the TCP-seq data. Read counts were converted to Reads per million (RPM). Distribution of Methionine specific elongator tRNA (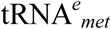) is also shown for the respective strains. **(B)** An outline of the methodology used to capture translation initiation and elongation complexes on an mRNA harbouring AGG start codon. **(C)-(D)** Distribution of the AGG-GFP mRNA derived FPs from the 30S (C) & 70S fractions (D) of Wt, *ΔrbfA* and *ΔksgA*. The FP counts are shown along the entire length of the gene encoding GFP.

Intrigued by the altered ribosome occupancy of the translation initiation complex formed by premature ribosomes bound to mRNA containing cognate start (AUG) codon (Fig. 4), we asked whether the identity of the cognate start codon has any role in limiting the progression of premature ribosomes towards elongation stage. To address this, we studied the distribution of 30S and 70S specific FPs from AGG-*gfp* (Fig. 5B). We reasoned that the distribution of FPs should mirror the observations from GFP and bgal expression studies (Fig. 1 & 2). In line with our conjecture, FPs derived from Wt 30S were densely populated around the start codon region (Fig. 5C), whereas 30S from *ΔrbfA* and *ΔksgA* showed reduced distribution of FPs around the start codon. This hints at the possibility that 30S from Wt stalls at the initiation site longer than the pre-30S from *ΔrbfA* and *ΔksgA*. This further suggests that unlike the pre-30S-PIC/IC from *ΔrbfA* and *ΔksgA*, 30S-PIC/IC from Wt does not efficiently progress towards the formation of 70S-IC if the start codon is AGG. Next, in order to understand how this 30S-PIC/IC transitioned into 70S-IC or an elongation complex, we analysed the distribution of FPs from the 70S fraction. Interestingly, we noted that despite the significant distribution of FPs around the initiation site from the Wt 30S and 70S fraction, their distribution throughout the length of GFP is abysmal, suggesting that the AGG-*gfp* is poorly translated (Fig 5C & 5D). Strikingly, 70S FPs from *ΔrbfA* were evenly distributed in higher proportion than those from *ΔksgA* and Wt, suggesting that premature ribosomes from *ΔrbfA* undergo rapid conversion of pre-30S-PIC/IC to 70S-IC, leading to efficient translation of AGG-*gfp*. This also agrees with the observations from GFP and bgal expression studies (Fig. 1 & 2).

### Hotspots for maintaining translation fidelity are compromised in premature ribosomes

Our experiments clearly outline that although premature, these ribosomes possess essential hotspots to engage with the translational machinery. How the 30S-PIC engaging the cognate or non-cognate initiation codon bypassed the scrutiny during the translation initiation intrigued us. To understand this, we analysed the structures of ribosome in complex with Initiation factors or RAFs (Boehringer et al., 2012; Hussain et al., 2016; Razi et al., 2017). Taking cues from our analyses and previous observations, we found a significant overlap in the binding site of Initiation factors (IFs) and late stage assembly factors on the 30S subunit. We hypothesized that the structural distortions in the premature ribosomes might have tempered affinities for the IFs. Such premature ribosomes, when entering translation with either cognate or non-cognate initiation signal could bypass the fidelity checkpoints posed by IFs, thus leading to an inaccurate translation event. To test this conjecture *in vivo*, we simply elevated the cellular concentration of all three IFs individually by ectopic expression, expecting that an increased cellular concentration of IFs may restore the skewed mass balance ratio between IFs and premature 30S subunits. IRI values were calculated as a ratio between GFP fluorescence from cells with elevated IF concentration to that of cells with basal concentration of IF. An IRI value ∼1 would mean that the expression levels of GFP were unaffected upon overexpressing IFs, whereas IRI < 1 would indicate repression of translation upon overproduction of the respective IF. The IRI values were measured for constructs with either AUG or AGG start codon individually. In order to achieve this, IF production was triggered when cells just entered the log phase (OD_600_ ∼ 0.2–0.3) followed by induction of GFP production at mid-log phase (OD_600_ ∼ 0.6), thus allowing for cellular levels of the respective IFs to be elevated before *gfp* expression could initiate (*vide*. Methods).

Elevated levels of IF-1 had no significant effects on expression of GFP from AGG or AUG start codons for Wt or *ΔksgA* (Fig. 6A). In *ΔrbfA*, IF-1 specifically suppressed expression from AGG (Fig. 6A). Unusually, an indiscriminate repression of GFP production from both AGG and AUG start codons was observed when cellular level of IF-2 was elevated (Fig. 6B). It would seem plausible that overexpression of IF-2 somehow inhibited translation. Strikingly, raising IF-3 level specifically inhibited expression from AGG start codon (Fig. 6C). However unlike IF-1, IF-3 mediated inhibition of translation from AGG start codon is equally strong even in case of Wt or assembly deficient strains, highlighting the role of IF-3 in discriminating the CO-AC interactions on the 30S-PIC/IC. Collectively, these observations reaffirm our hypothesis that the restoration of the pre-30S-IF mass balance may avert the participation of premature ribosomal particles in translation. To further probe the IF mediated moderation of the ribosome, we studied the polysome profiles from Wt, *ΔksgA* and *ΔrbfA* containing elevated concentrations of the respective IFs (Fig. 6D-6F). In line with our expectations, complementation of RbfA and KsgA restored the distorted polysomes profiles in the respective null mutants (inset in Fig. 6E & 6F), suggesting efficient rescue from the assembly defects. Further, except in *ΔrbfA*, elevated level of IF-1 did not seem to alter the distribution of ribosomes in Wt and *ΔksgA* and in comparison to the basal level of IF1 in these strains (Fig. 6D-F). However, overexpression of IF-2 led to a stabilizing effect on the 70S particles that seemed drastic for *ΔrbfA* (Fig. 6E, compare *ΔrbfA* with *ΔrbfA* + IF-2) and subtler for Wt and *ΔksgA* (Fig. 6D & 6F). On the contrary, elevated levels of IF-3 displayed a pronounced subunit dissociation activity, where a decrease in the 70S population was accompanied by a concomitant increase in the 50S and 30S populations (Fig. 6D-6F). Similar to IF-2, the effect of IF-3 overexpression was more prominent in *ΔrbfA* as compared to Wt and *ΔksgA* (Fig. 6D-6F, compare *ΔrbfA* with *ΔrbfA* + IF-3).

**Figure 6:**
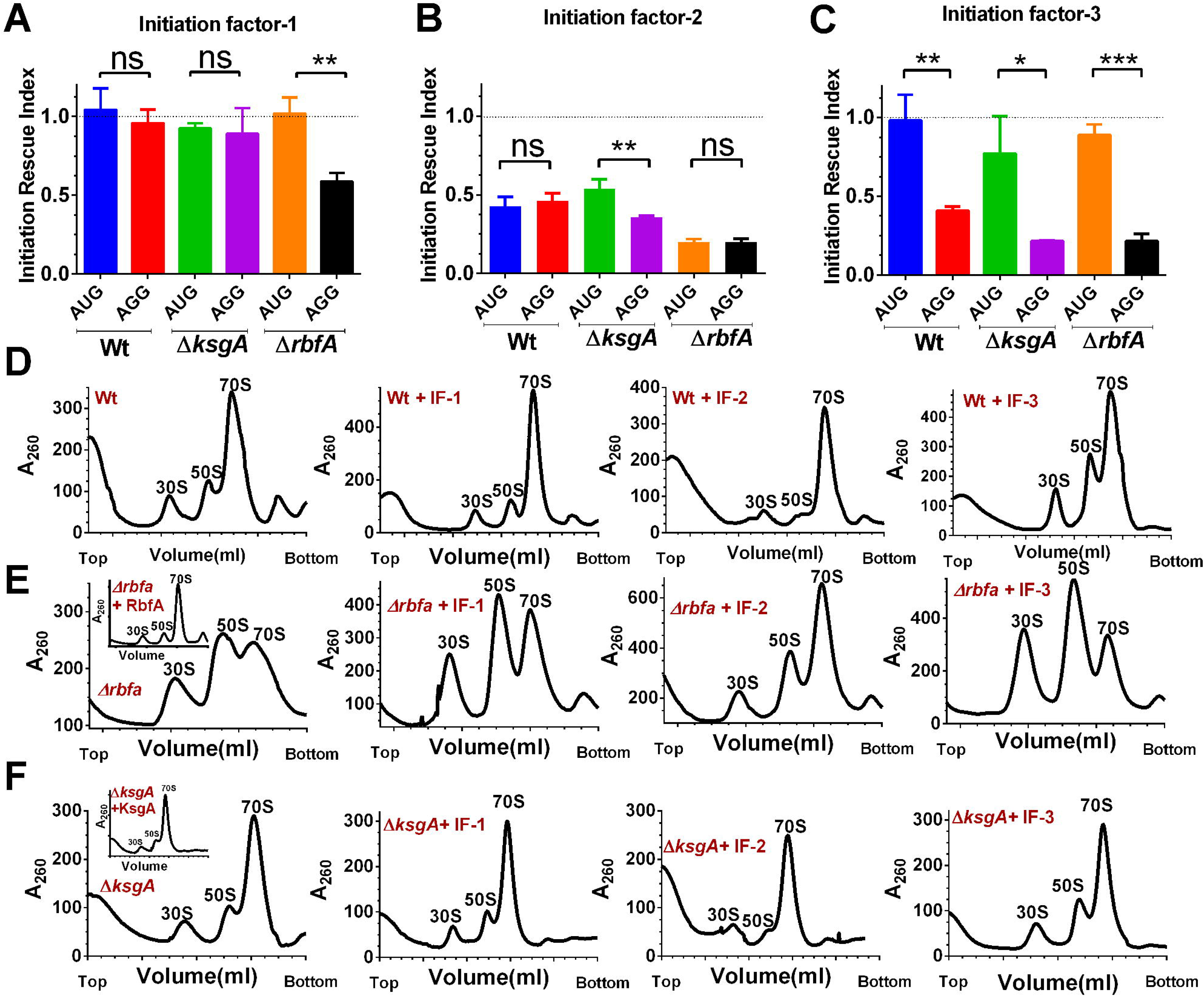
Elevated levels of initiation factors avert premature ribosomes from engaging in translation. **(A)** IRI measurements for Wt, *ΔrbfA* and *ΔksgA* from a *gfp* encoded mRNA with AUG or AGG start codon in presence of elevated cellular concentrations of IF-1. The statistical significance was tested using a paired t-test analysis with a 95% confidence interval. **(B)** IRI measurements for Wt, *ΔrbfA* and *ΔksgA* from a *gfp* encoded mRNA with AUG or AGG start codon in presence of elevated cellular concentrations of IF-2. **(C)** IRI measurements for Wt, *ΔrbfA* and *ΔksgA* from a *gfp* encoded mRNA with AUG or AGG start codon in presence of elevated cellular concentrations of IF-3. **(D)** Representative polysome profiles for Wt cells carrying an empty p15A vector (only Wt) or p15A vector carrying genes encoding IF-1, IF-2 or IF-3. The profiles were drawn from mid-log phase cells overexpressing the respective initiation factors. **(E)–(F)** Representative polysome profiles for *ΔrbfA* or *ΔksgA* cells carrying an empty p15A vector (only *ΔrbfA* or *ΔksgA)*. The inset shows the profile for the respective null mutants carrying p15A vector carrying a gene encoding RbfA or KsgA. Similar profiles are also shown for the respective null mutants carrying the p15A vector encoding IF-1, IF-2 or IF-3.

In parallel, we also checked if ectopic expression of IFs improved cellular fitness, which could in turn lead to decrease in error-prone translation. To test this, Wt, *ΔrbfA* and *ΔksgA* cells transformed with plasmids carrying genes encoding IFs were tested for their growth characteristics after inducing the expression of the respective gene encoding IF. For the purpose of uniformity, OD_600_ of all cultures was normalized at the time of starting the experiment and IF expression was also triggered at the same time (time = 0) (*vide* methods). The absence of a growth advantage (Fig. S3) indicated that the rescue is only a result of the translation quality control and not due to any increase in the overall pool of premature ribosomes. As observed, overexpression of IF-1 or IF-3 did not improve cellular fitness in any manner but a radical effect of IF-2 overexpression was observed for all the three strains as they failed to grow even after prolonged incubation (Fig. S3). These observations suggest a central role of IF-2 in regulation of translation (Milon et al., 2006), that may be very sensitive to cellular concentrations of IF-2.

### Repression of translation from mRNA with non-cognate start codon is at the initiation stage

It is evident from the above experiments that IFs pre-empt premature ribosomes with assembly defects from participating in translation. However, in order to understand the mechanism of this inhibition we studied translation kinetics in strains with elevated level of IFs. To perform this, the production of IF was triggered before measuring bgal production kinetics. Elevated IF-1 level displayed appreciable disruption of translation initiation events only from AGG start codon in *ΔrbfA* (Fig. S4A & S4D; Table 3). This was in line with the GFP repression from AGG start codon for *ΔrbfA* during the steady state measurement of GFP production (Fig. 6A). However, in case of IF-2 overexpression, the initiation events dampened for AUG start codon and fell abysmally for AGG start codon (Fig. S4B & S4E; Table 3). A completely different scenario unfolded when the cellular concentration of IF-3 was increased. Here, the initiation events were not repressed appreciably for AUG start codon but a calibrated decrease was seen selectively for AGG start codon (Fig. S4C & S4F; Table 3). This IF-3 mediated progressive inhibition of translation from non-canonical initiation signals again reinforces the positive effects of restoration of the mass balance between IFs and premature subunits

**Table 3:** Translation initiation rates (IR) measured from Wt, *ΔksgA*, and *ΔrbfA* cells after elevating cellular concentrations of IFs. Translation initiation from the respective start codons is shown for cells induced (elevated) or uninduced (basal) for IF overproduction. The IR values are derived as the slope of the curve marking accumulation of bgal over time (units: Miller units^1/2^/sec).

| Strains | Start codon | IF-1 basal | IF-1 elevated | IF-2 basal | IF-2 elevated | IF-3 basal | IF-3 elevated |
| --- | --- | --- | --- | --- | --- | --- | --- |
| **Wt** | AUG | 0.09065 ± 0.003 | 0.09714 ± 0.004 | 0.1014 ± 0.002 | 0.08108 ± 0.003 | 0.1113 ± 0.005 | 0.09598 ± 0.005 |
| **Wt** | AGG | 0.1115 ± 0.006 | 0.1156 ± 0.006 | 0.07005 ± 0.003 | 0.03771 ± 0.000 | 0.04192 ± 0.000 | 0.02794 ± 0.001 |
| *ΔksgA* | AUG | 0.07674 ± 0.004 | 0.05927 ± 0.003 | 0.08411 ± 0.002 | 0.04730 ± 0.002 | 0.1213 ± 0.006 | 0.1044 ± 0.003 |
| *ΔksgA* | AGG | 0.1001 ± 0.006 | 0.09060 ± 0.011 | 0.08254 ± 0.004 | 0.01587 ± 0.001 | 0.07259 ± 0.002 | 0.05026 ± 0.003 |
| *ΔrbfA* | AUG | 0.07567 ± 0.003 | 0.06036 ± 0.002 | 0.07105 ± 0.003 | 0.04475 ± 0.003 | 0.06621 ± 0.002 | 0.06027 ± 0.001 |
| *ΔrbfA* | AGG | 0.1465 ± 0.009 | 0.1017 ± 0.003 | 0.09060 ± 0.002 | 0.01766 ± 0.001 | 0.08442 ± 0.003 | 0.04989 ± 0.002 |

## Discussion

Genetic disruptions of ribosome assembly (Dragon et al., 2002; Ganapathi et al., 2007; Sun and Woolford, 1994) and production of misfolded proteins by a compromised translational machinery have been associated with cell death and neurodegeneration (Lee et al., 2006). In order to counter this, eukaryotic systems deploy proactive mechanisms to avert the entry of premature ribosomes into the translation cycle (Strunk et al., 2012). However, parallel prokaryotic mechanisms that may ensure entry of only mature ribosomes into translation still elude discovery. Interestingly in *E. coli*, late stages of 30S subunit maturation seem to be closely coupled with translation initiation underscoring the intimate link between assembly and translation initiation (Shetty and Varshney, 2016). Our data suggests that quality control checkpoints are also present in bacterial cells and that premature ribosomes produced in the absence of RAFs successfully evade these checkpoints. Such premature ribosomes possess suboptimal fidelity to recognize the cognate (AUG) start codon (Fig. 1). Consistent with previous report of mistranslation events in *ΔksgA* (Connolly and Culver, 2013), our observations for *ΔrbfA, ΔksgA* and *ΔlepA* also reiterate a direct correlation between assembly defects and impaired translational machinery. It is important to reflect upon, whether the mistranslation events observed here arise from CO-AC interactions between non-cognate mRNA and 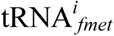 or incorporation of tRNA*^e^* into the pre-30S-ICs. Our observations of significant 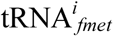 localization in 70S particles from *ΔrbfA* and *ΔksgA* (Fig. 5A) and previous reports of N-terminal sequencing of mistranslated products (O’Connor et al., 1997) suggest that such misinitiation events arise due to interactions between the 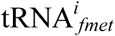 and non-AUG start codons. This further attributes mistranslation to a failure in discriminating non-canonical CO-AC interactions during formation of the pre-30S-IC.

What is lacking in pre-30S so that it looses the ability to discriminate non-canonical CO-AC interactions during initiation? During assembly, RbfA, which is a cold shock protein, is proposed to assist in formation of helix h1 (Dammel and Noller, 1995) and later to reorient h44 and h45 that form the 3’minor domain of 16S rRNA. Further, during these late stages of assembly, KsgA modifies two consecutive residues on the h45 region (Helser et al., 1972; Xu et al., 2008). Incidentally, mutations in h44 also promote initiation from non-AUG start codon by altering the movement of the 3’ major domain that forms the 30S head region (Qin and Fredrick, 2009). All these findings, hint at an essential need for efficient maturation of the 16S 3’ domain for the decoding step. In this study, we find that recognition of canonical start signals seems to remain unaffected but there is a gradual increase in the propensity to recognize near-cognate (CUG and AUA) or a non-cognate (AGG) start codon for both *ΔksgA* and *ΔrbfA* (Fig. 1). Elevated expression from AGG start codon is particularly noteworthy given the marked conservation of U among other near-cognate start codons (i.e., G<u>U</u>G, U<u>U</u>G, A<u>U</u>U, A<u>U</u>C, and A<u>U</u>A) and indicates severely compromised decoding capabilities at the 30S P-site. However, milder defects in recognizing frameshifts and stop codons (Fig. 1G) indicate a limited effect of assembly defects on decoding at the A-site too. Further, our experiments to gauge the mechanistic implications of impaired P-site decoding also reveal wide spread effects on protein production due to a decrease in rate of translation initiation and peptide bond synthesis (Fig. 2C, 2D & 2F). However, it is not possible to infer from the current data, if this decrease in the peptide chain elongation rate is solely due to defects in the P-site or other structural deformities that decrease the speed of the translocation step (Shi et al., 2009), thus slowing down the overall process of translation. It also seems plausible to extend that as defects in assembly exacerbate, defects in codon recognition and translation impairment also aggravate (Fig. 3C, 3D & 4). However, it is evidently clear that these defects are not an outcome of translation cessation in RAF null mutants but rather a bona fide compromise in the fidelity (Fig. 2F).

Led by the cue that the pre-30S harbours defect that has compromised the fidelity of decoding, we asked what aspect of initiation falters. During the initiation stage, the 30S binds the Initiation factors (IF-1, IF-2 and IF-3), the mRNA and the 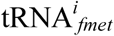 in a poorly defined order to form the 30S-PIC, wherein the formation of cognate CO-AC interaction is not yet complete. A large-scale conformational rearrangements within the 30S-PIC guided by the Initiation factors ensure the proof reading of cognate CO-AC interaction and the transition towards the 30S-IC, which subsequently docks with the 50S to form the 70S-IC that is competent to enter the elongation cycle (Hussain et al., 2016; Milon et al., 2012). Here, the transition of 30S-PIC to 30S-IC is governed by the Initiation factors and this stage represents the first checkpoint to ensure the translational fidelity. Our analysis shows that it is the same pathway that is manned by Initiation factors to ensure the cognate CO-AC interaction at the P-site goes awry when 30S harbours assembly defects. It is possible to discern two aspects during the transition from 30S-PIC to 30S-IC that seem to be critical in determining the fate of bona fide initiation. These are (i) the formation of cognate CO-AC interaction and (ii) the proof reading of cognate CO-AC interaction by Initiation factors (especially IF3). In the case of Pre-30S, the Initiation factors bind weakly and therefore this impacts the proof reading of cognate CO-AC interaction (Fig. 6, S4 & 7; Table 3). This is further exacerbated when the pairing is formed between AGG and 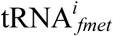, wherein both the formation of cognate CO-AC interaction and its subsequent proof reading by the Initiation factors are disrupted. Unlike the Wt, the altered ribosome occupancy as well as the preferential enrichment of 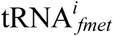 with 70S particles in the case of *ΔksgA* and *ΔrbfA* suggest rapid transition of 30S-PIC/IC to 70S-IC possibly by evading the translation quality control checkpoint (Fig. 4 & 5A). This is bolstered by the robust expression of AGG-*gfp* using pre-30S (Fig. 1E, 1F, 2E, 2F, 5C & 5D). These notions have been reaffirmed by the initiation rescue observed when cellular concentrations of IFs were increased to compensate for a feeble pre-30S-IF binding due to structural defects in the pre-30S (Fig.6A-6C). It is however noteworthy, that IF-1 and IF-3 seem to specifically inhibit translation initiation (Fig. S4) while the reason for non-specific inhibition of translation initiation after IF-2 overexpression is not clear (Fig. 6, S4B & S4E). One plausible scenario for this could be that the elevated IF-2 levels are not appropriately accounted for by the concomitant rise in the levels of 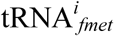, leading to suboptimal charging of IF-2 with 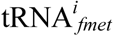. This may give rise to vacant IF-2 binding to 30S and stalling of initiation. IF-3 is already known to discriminate the CO-AC interaction during the formation of the 30S-PIC and its subsequent conversion to 30S-IC by a series of precisely controlled structural rearrangements (Hussain et al., 2016; Milon et al., 2008; Milon et al., 2012). Its subunit anti-association activity (Fig. 6D-6F) also seems to play critical roles in ensuring fidelity of initiation by rapid ejection of non-canonical mRNA-tRNA in the P-site, which could be abolished in case of pre-30S (Petrelli et al., 2001). However, the mechanism of IF-1 mediated rescue is subtler (Fig. 6D-6F) as it is thought to enhance the function of IF-2 and IF-3 (Hussain et al., 2016). Thus it seems possible to derive from these findings that a speedy conversion of initiating pre-30S particles to elongating 70S particles takes place by skipping the IF mediated quality control of protein synthesis (Fig. 7). Further structural and biochemical evidence would be needed to understand the nature of these structural defects to fully understand the factors that contribute to a compromised decoding fidelity in premature subunits.

**Figure 7:**
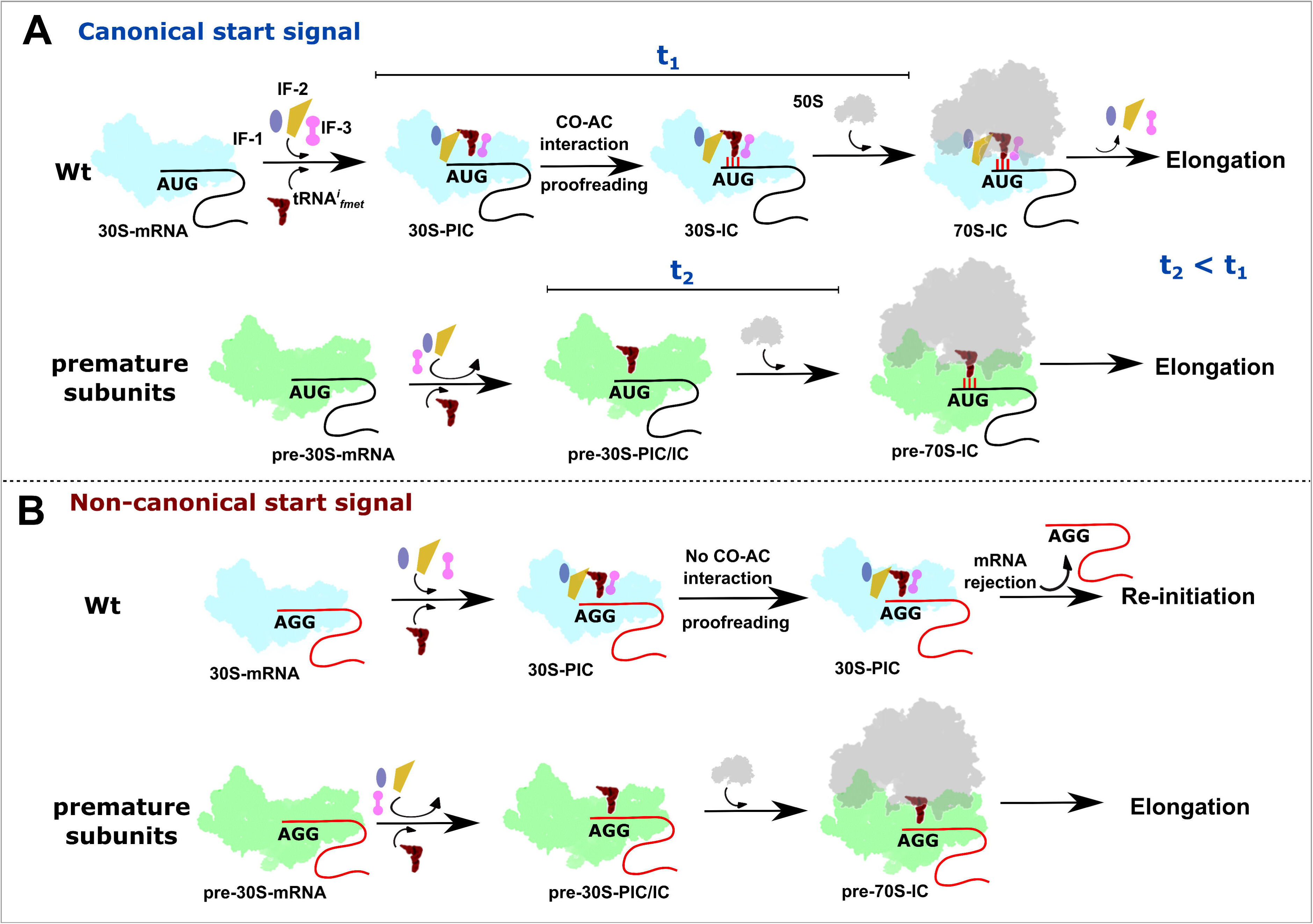
A schematic representation of the plausible events leading to the participation of premature subunits in translation. **(A)** Initiation factors and 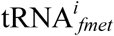 are recruited to mature 30S subunits complexed with mRNA harbouring canonical start signal to form accurate initiation intermediates. These intermediates undergo meticulous kinetic proofreading checkpoints posed by initiation factors and only then are allowed to enter translation. On the contrary, premature subunits with structural deformities possess lower binding affinities for initiation factors thus readily bind to 50S particles to initiate translation from canonical start signals. This further suggests that the time for forming cognate codon-anticodon pairing and subsequent proofreading by IFs is faster for premature subunits (t2) than for mature subunits (t1). (B) Mature 30S particles can form pre-initiation complexes on non-canonical signals, but proofreading of the impaired codon-anticodon interaction by initiation factors preempts their transition into initiation complexes and further translation. However, due to the weak binding of Initiation factors, premature 30S particles elude such proofreading checkpoints and undergo rapid transition to form 70S, which initiates protein synthesis from non-canonical initiation signals thus displaying compromised fidelity of translation initiation.

## Concluding remarks

In bacteria, it appears that the translation Initiation factors act as a single window quality control checkpoint to proofread not only the fidelity of the genetic message during translational cycle but also to test drive and validate the competence of nascent ribosomes. In this context, it is tempting to speculate that the universality of AUG as an initiation codon perhaps stems from the fact that the initiation factors, especially IF3, is evolved and attuned to recognize AUG than the non-AUG. This inter-dependency is also a cost-cutting measure to optimize the energy reserves while at the same time tightening the quality control.

## Data and Software Availability

Sequencing data are deposited in the Gene Expression Omnibus under accession number GEO: GSE122870

## Acknowledgements

This work was supported by grants from Department of Biotechnology (DBT) [BT/PR15925/NER/95/141/2015, BT/08/IYBA/2014/05, BT/406/NE/UEXCEL/2013, BT/PR5511/MED/29/631/2012 and BT/341/NE/TBP/2012] and Science and Engineering Research Board (SERB) [YSS/2014/000286]. The pBAD Strep TEV LIC (8R) cloning vector (Addgene plasmid # 37506) and the pET His6 GFP TEV (1GFP) LIC cloning vector (Addgene plasmid # 29663) were a gift from Scott Gradia. The vector pdCas9-bacteria was a gift from Stanley Qi (Addgene plasmid # 44249). We acknowledge the geniality of the aforementioned scientists for sharing their plasmids. We also acknowledge the NGS services of Genotypic Technologies Pvt. Ltd., Bengaluru, India. The authors also thank Sumit Kinger and Kiran Dhobale for the technical assistance and are also grateful to all members of the MAB lab for their support and critical comments on the manuscript.

## Author contributions

H.S. and B.A. conceived the problem; H.S. designed assays and performed all experiments; B.A processed and analysed the Deep sequencing data; H.S. and B.A. analysed the remaining data and wrote the manuscript; B.A. arranged for funds and oversaw the project.

## Declaration of Interests

The authors declare no conflict of interest

## Experimental procedures

### Creation of strains and plasmids

The Keio collection parent strain BW25113 referred as Wt was used as the parental strain for all genetic manipulations and reference measurements. Null mutant for LepA was procured from the Coli genetic stock centre (CGSC). Additionally, null mutants for RbfA, RsgA and KsgA were created using λ Red recombineering method (Datsenko and Wanner, 2000). Gene encoding GFP was amplified from the vector 1GFP (Addgene #29663) with four different start codons i.e. AUG, CUG, AUA and AGG. Additionally, frameshift mutations (+1 base and −1 base) were introduced at codon 7 of the GFP construct using oligonucleotides. Similarly, codon 7 and 8 were replaced with UAG and UAA codons. The modified GFP constructs were individually cloned into vector 8R (Addgene # 37506) using Xba1/Nhe1 and BamH1 specific restriction sites. Similarly, the constructs used for studying translation kinetics were created by amplifying the gene encoding β-galactosidase (bgal) from *E. coli* B cells. Modifications were also introduced into this gene by incorporating different start codons (O’Connor, 1997; O’Connor et al., 2004). The amplified cassettes were cloned into vector pQE2 using XhoI/HindIII sites. In order to complement assembly factors (RAFs) or initiation factors (IFs), respective genes were amplified from Wt cells and cloned into a p15A vector backbone under an Anhydro-tetracycline (Atc) inducible promoter.

### Growth analysis

Primary inoculums were always prepared by growing freshly transformed colonies in LB medium with respective antibiotics at 37 °C with shaking at 180 rpm. The next day, OD_600_ of primary cultures was normalized. These cultures were then diluted into 1 ml fresh LB medium with appropriate antibiotics in a 24 well plate followed by incubation at 37 °C with regular shaking. OD_600_ measurements were taken in Tecan infinite M200 multimode plate reader at every 30-minute interval until cultures reached saturation. Additionally, in order to study growth in presence of elevated levels of IFs or RAFs, LB medium was supplemented with 50 ng/ml Atc at the time of inoculation to trigger protein production. The experiment was repeated at least three times to derive the average growth curves.

### GFP fluorescence measurements and calculation of indices

Wt or null mutant was freshly transformed using the desirable plasmid carrying the gene encoding GFP before initiating the experiments. Colonies were picked in fresh LB medium supplemented with 100 μg/ml ampicillin (Amp) and allowed to grow at 37 °C and 180 rpm till OD_600_ reached 0.6. At this point, *gfp* expression was induced by adding 2 mM arabinose and cells were allowed to grow for another 3 hrs. Upon completion, cells were harvested and lysed in Buffer G (20mM Tris-HCl (pH 7.5 at 25 °C), 500 mM NaCl, 1 mM EDTA, 1 mM PMSF, 6 mM β-ME) supplemented with 1X CelLytic B solution (Sigma Aldrich). The lysates were normalized for total protein content and were taken for fluorescence measurements. All fluorescence spectrums were generated by exciting at 488 nm and scanned for emission from 500–600 nm, with averaging over three scans after baseline correction in a FluoroMax-4 spectrofluorometer (Horiba Scientific, Edison, NJ). The slit width used for excitation and emission was 2 nm and 7 nm, respectively. All fluorescence experiments were performed with five independent trials. For measurement of Initiation Error Index, GFP emission from null mutants was compared with that of Wt cells. Similarly, Rescue Index was calculated as the ratio of GFP fluorescence after complementing the null mutant with respect to Wt. Briefly, cells were co-transformed using plasmids carrying genes coding for GFP and assembly or initiation factor, respectively. The cells were grown on a dual selection marker at 37 °C and induced with Atc to trigger production of the RAF or IF when OD_600_ reached 0.3. Subsequently, GFP production was induced with 2 mM arabinose when OD_600_ of the Atc induced cells reached 0.6. Further, the cells were processed for fluorescence measurements as described above. The indices were defined as following:

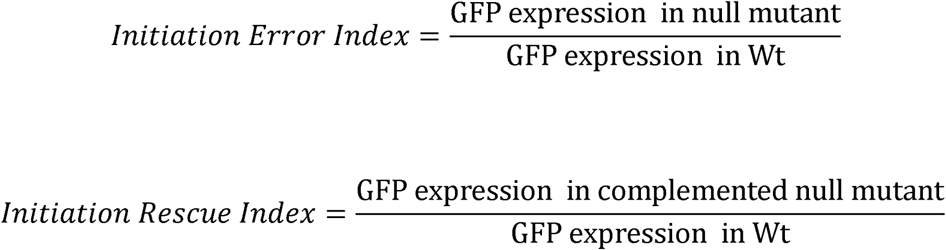

### Determination of initiation and elongation rates from β-galactosidase (bgal) production kinetics

In order to calculate the translation initiation and elongation rates, the bgal production assay (Dalbow and Bremer, 1975; Dalbow and Young, 1975; Miller, 1992) was performed with minor modifications. Briefly, Wt or null mutant was transformed using respective constructs that encoded bgal (variants containing different start codons: pAUG-bgal or pAGG-bgal). Overnight grown cultures were diluted into fresh 5 ml LB medium supplemented with 100 μg/ml Amp and allowed to grow at 37 °C and 180 rpm till OD_600_ reached 0.6. These cultures were then shifted to a water bath at 37 °C or 25 °C (RT). Bgal production was induced with 1 mM IPTG followed by instant mixing for 5 secs. Following this, aliquots of 200 μL each were taken every 30 secs for the assays done at 37°C and every 120 secs for assays done at RT. The aliquots were added to tubes containing 300 μL of Z-buffer (60 mM Na_2_HPO_4_·2 H_2_O, 40 mM NaH_2_PO_4_· H_2_O (pH 7.0 at 25 °C), 10 mM KCl, 1 mM MgCl_2_, 5 mM β-ME), 20 μL of chloroform, and 10 μL of 0.1% SDS. The tubes were then instantly vortexed for 20 secs and transferred to ice. At the end, all the tubes were incubated for 5 mins on ice and then for 5 mins at RT after which, 500 μL of 1 mg/ml ONPG (o-nitro-phenyl galactopyranoside) was added to the reaction. ONPG hydrolysis for cells with pAUG-bgal plasmid was allowed to proceed for 15 mins at RT. Similarly, for cells with pAGG-bgal the reaction was allowed to proceed for 1 hour at RT. After completion, the reaction was stopped by adding 500 μL of 1 M Na_2_CO_3_. Colour was allowed to develop for 15 mins and then 300 μL of the reaction mix was transferred to 96 well plates. Absorbance was recorded at 420 nm and 550 nm using a Tecan infinite M200 multimode plate reader. The readings were used to derive the plot representing residual enzymatic activity for calculating elongation rates and rates of initiation (Schleif et al., 1973). Length of the protein (1,024 amino acids) divided by the lag time for the appearance of enzymatic activity after induction was used to calculate the elongation rates. The square root of the residual enzyme activity √(E_t_-E_0_), where E_t_ signifies the miller units at time t and E_0_ signifies the miller units recorded at t=0 was plotted against time (t) to derive a linear plot for calculating the rate of translation initiation.

### Polysome profiling

Wt, *ΔrbfA* or *ΔksgA* cells were grown overnight in LB medium. For complementation studies, these cells were transformed with a plasmid carrying the respective *rbfA*, *ksgA, infA*, *infB* or *infC* and were grown overnight in LB medium supplemented with 25 μg/ml Chloramphenicol (Cmp). The next day, cultures were diluted in 100 ml fresh LB medium supplemented with Cmp and grown at 37 °C with vigorous shaking. After OD_600_ reached 0.3, the protein production was induced with Atc (50 ng/ml) and allowed to grow till the OD_600_ of the cultures was 0.6. At this point, the culture was chilled rapidly on ice with addition of ice cubes and 200 μg/ml Cmp followed by incubation for 15 mins. The cells were harvested by centrifugation at 4 °C, 8000xg for 10 mins. Cells were then washed once with Buffer CL (20 mM Tris-HCl (pH 7.6 at 25 °C), 150 mM NH_4_Cl, 10.5 mM Mg(OAc)_2_, 0.5 mM EDTA and 6 mM β-ME) and later re-suspended in 1 ml buffer CL supplemented with 1 mg/ml Lysozyme, 1 mM PMSF, 0.5x CellLyticB reagent followed by incubation on ice for 1 hour (hr). The cells were then ruptured by 5 cycles of freeze-thaw using liquid nitrogen. The lysates were clarified by centrifugation at 30,000xg followed by A_260_ (Absorbance at 260 nm) quantification in Implen Nanospectrophotometer. For ultracentrifugation, 300μL of 10 A_260_ was loaded on a 10–50% sucrose gradient prepared in buffer CL and spun at 102,000xg in rotor TH-641 (Thermo-Sorvall WX^+^ ultracentrifuge) for 16 hrs. Subsequently, fractionation was performed using a syringe pump connected to Akta Pure system (GE healthcare).

### Growth, Cross-linking and cell lysis conditions for TCP-seq

Preparation of ribosome and initiation complex footprints were done essentially according to the original protocol (Shirokikh et al., 2017) with minor modification as required for bacterial cells. Overnight cultures of Wt, *ΔrbfA* or *ΔksgA* cells carrying the plasmids pAGG-GFP-8R were diluted into 2 litres of fresh LB medium supplemented with 100 μg/ml Amp and incubated at 25 °C with shaking at 180 rpm. Minimal *gfp* expression was induced using 50 μM arabinose and growth was allowed to proceed till OD_600_ reached 0.4. At this point, cells were rapidly chilled by adding ice cubes accompanied by addition of 150 ml freshly prepared 30% formaldehyde. Cultures were mixed vigorously and placed on ice for 15 mins followed by addition of 200 ml 2.5 M glycine. Following this, cells were harvested by centrifugation at 8000xg for 10 mins. Pellets were washed with 40 ml of Buffer TW (20 mM HEPES-KOH (pH 7.4 at 25 °C), 100 mM KCl and 2 mM MgCl_2_) followed by resuspension in 0.4 ml of buffer TL (20 mM HEPES-KOH (pH 7.4 at 25 °C), 100 mM KCl, 2.5 mM MgCl_2_ and 0.5 mM EDTA) supplemented with 2 mg/ml lysozyme, 6 mM β-ME, 1 Unit/μl RNaseOUT inhibitor (ThermoFisher Scientific), 40 Units DNase and 1 mM PMSF. Cells were placed on ice for 1 hr followed by rupturing using 5 cycles of freeze-thaw in liquid nitrogen. Cell lysates was clarified by centrifugation at 30,000xg for 30 mins at 4 °C.

### Isolation of mRNA bound fractions and MNase treatment

Clarified cell lysate was layered onto a 10–20% sucrose gradient prepared in Buffer TP (50 mM Tris-HCl (pH 7.0 at 25 °C), 50 mM NH_4_Cl, 4.5 mM MgCl_2_, 0.5 mM EDTA, 6 mM β-ME) and centrifuged at 52,000 rpm for 80 mins at 4 °C using a TLS-55 rotor in an Optima TLX 120 ultracentrifuge (Beckman-Coulter). Ribosome pellets derived after centrifugation were resuspended in 1 ml of buffer TL supplemented with 1 Unit/μl RNaseOUT and 1 mM PMSF. These ribosomes were then treated with 30 Units of MNase per 1 A_260_ of ribosomes for 1 hr at RT. The reaction was stopped by addition of 2 mM EGTA and the nuclease treated complexes was immediately loaded onto a 10%-40% sucrose gradient prepared with buffer TP. Samples were spun at 40,000 rpm for 4 hrs at 4 °C using a TLS-55 rotor. Upon completion, fractions of 100 μl each were collected manually and later processed for footprint isolation.

### De-crosslinking and footprint isolation

Fractionated 30S or 70S samples were supplemented with 1% w/v sodium dodecyl sulphate (SDS), 10 mM EDTA, 10 mM Tris-HCl (pH 7.4 at 25 °C), 10 mM glycine and 1 mg/ml Proteinase K (Sigma-Aldrich). The samples were incubated at 50 °C for 30 mins with regular mixing. Following this, equal volume of acid phenol: chloroform (5: 1) was added to this mix and incubated at 65 °C for another 45 mins with regular vortexing. The sample was then centrifuged at 16,000xg for 30 mins at 4 °C followed by separation of the aqueous phase. Equal volume of chloroform was then added to the aqueous phase and centrifuged at 16,000xg for 30 mins at 4 °C. The aqueous phase was again separated in a fresh tube and supplemented with 0.1 volume of 3 M sodium acetate (pH 5.2 at 25 °C), 1 mg/ml glycol blue (Ambion) and 2.5 volumes of chilled ethanol. Precipitation was done overnight at −20 °C following which the sample was centrifuged at 16,000xg for 45 mins at 4 °C. RNA pellets were washed twice with chilled 70% ethanol followed by air drying. Finally, the pellets were resuspended in nuclease free water and checked for integrity. Footprints corresponding to 30S or 70S fractions were size selected from 10–80 bases by gel elution using denaturing Polyacrylamide gel electrophoresis (PAGE) in Protean ii XL setup (Bio-Rad). The extracted RNA was end repaired using T4 Poly nucleotide kinase (PNK) and purified by ethanol precipitation.

### Library preparation and sequencing

Library preparation and sequencing was outsourced to Genotypic Technology Pvt. Ltd., Bengaluru, India. Libraries were prepared strictly according to the manufacturers recommendations using the TruSeq Small RNA Sample Preparation kit (Illumina, U.S.A). Briefly, 50 ng of RNA was used as starting material for ligation of 3’ and 5’ adapters. Specific index sequence was added to each sample for identification during sequencing. The Illumina Universal Adapter used in the study was:

AATGATACGGCGACCACCGAGATCTACACGTTCAGAGTTCTACAGTCCGA and the Index Adapter was:

CAAGCAGAAGACGGCATACGAGAT[INDEX]GTGACTGGAGTTCCTTGGCACCCGAGA ATTCC.

Adapter ligated fragments were reverse transcribed with Superscript III Reverse transcriptase (Invitrogen). The cDNA was enriched and barcoded by 15 cycles of PCR amplification and the amplified library was size selected using denaturing PAGE. The library was size selected in the range of 140 bp– 210 bp followed by overnight gel elution and finally resuspended in nuclease free water. Illumina compatible sequencing library was initially quantified by Qubit fluorimeter (Thermo Fisher Scientific) and its fragment size distribution was analysed on Agilent TapeStation. Sequencing was performed on the Illumina NextSeq 500 platform. The depth of sequencing for each sample ranges from 10 to 15 million reads.

### Analysis of sequencing data

The reads were subjected to several pre-processing steps described as follows. Firstly, reads with Phred score less than 20 were removed by utilizing fastq_quality_trimmer from the FASTX-toolkit-version-0.0.13. The remaining reads were trimmed for 3’-end adapter sequences [5’-TGGAATTCTCGGGTGCCAAGGAACTC-3’] using Cutadapt-1.5 (Martin, 2011). Following these, in order to filter and remove the reads derived from rRNA, tRNA, sRNA or any other non-coding RNA, reads were aligned using STAR-2.5.3 (Dobin et al., 2013) against non-coding RNAs from *E. coli* K-12 MG1655 genome (ftp://ftp.ncbi.nlm.nih.gov/genomes/refseq/bacteria/Escherichia_coli/reference/GCF_000005845.2_ASM584v2/GCF_000005845.2_ASM584v2_rna_from_genomic.fna). The unmatched reads were aligned against the *E. coli* K-12 MG1655 reference genome (NCBI: NC_000913.3;ftp://ftp.ncbi.nlm.nih.gov/genomes/refseq/bacteria/Escherichia_coli/reference/GCF_000005845.2_ASM584v2/GCF_000005845.2_ASM584v2_genomic.fna) using STAR-2.5.3 with the following parameters: spliced alignment turned off (--alignIntronMax 1), forced end to end alignment (--alignEndsType EndToEnd) and allowing 8 mismatches per 100 nt (--outFilterMismatchNoverLmax 0.08). Sam to Bam conversion was accomplished using Samtools-1.4.1 (Li et al., 2009). Only uniquely matching reads were considered for further analyses. Metagene analysis for mapping the reads to known START codon of 4257 CDS in *E. coli* was performed by employing custom written shell scripts utilizing Samtools-1.4.1. Plots were generated using ggplot2 implemented in R.

